# DOT1L Complex Regulates Transcriptional Initiation in Human Cells

**DOI:** 10.1101/2020.12.07.414722

**Authors:** Aiwei Wu, Junhong Zhi, Tian Tian, Lixue Chen, Ziling Liu, Lei Fu, Robert G. Roeder, Ming Yu

## Abstract

DOT1L, the only H3K79 methyltransferase in human cells and a homolog of the yeast Dot1, normally forms a complex with AF10, AF17 and ENL/AF9, is dysregulated in most of the cases of mixed lineage leukemia (MLL) and is believed to regulate transcriptional elongation without much evidence. Here we show that the depletion of DOT1L reduced the global occupancy without affecting the traveling ratio or the elongation rate of Pol II, suggesting it not a major elongation factor. An examination of general transcription factors (GTFs) binding revealed globally reduced TBP and TFIIA occupancies near promoters after DOT1L loss, pointing to a role in transcriptional initiation. Proteomic studies uncovered that DOT1L regulates transcriptional initiation likely by facilitating the recruitment of TFIID. Moreover, ENL, a DOT1L complex subunit with a known role in DOT1L recruitment, also regulates transcriptional initiation. Furthermore, DOT1L stimulates H2B monoubiquitination by limiting the recruitment of human SAGA complex, and the connection between Dot1/DOT1L and SAGA complex is conserved between yeast and human. These results advanced current understanding of roles of DOT1L complex in transcriptional regulation and MLL.

## INTRODUCTION

Transcription is the first step of gene expression, cell type-specific transcription is fundamental to the development of multicellular organisms, and mutations of transcription factors (TFs) are found in more than 50% of all cancer cases ^1^. Among the three eukaryotic RNA polymerases, i.e., Pol I, II and III, the regulation of Pol II transcription is the focus of research because of the large number of protein-coding genes with varying lengths and therefore the complexity of the underlying mechanisms. Transcription is divided into three stages, including initiation, elongation and termination. In the initiation of Pol II transcription, six general transcription factors (GTFs), i.e. TFIIA, B, D, E, F and H, form a preinitiation complex (PIC) with Pol II for the recognition of a transcriptional start site (TSS) and the creation and stabilization of a transcription bubble ^2^. TFIID contains TBP and 13 TBP-associated factors (TAFs), and is critical for the recognition of promoters and the subsequent PIC assembly ^3^. Interestingly, three of the TAFs (TAF9, 10 and 12) were also found to be parts of the Spt-Ada-Gcn5-acetyltransferase (SAGA) complex, which contains TBP loading, activator binding, acetyltransferase, and deubiquitinase modules ^4^. In the yeast *Saccharomyces cerevisiae,* the TBP loading function and therefore a role in transcriptional initiation of the SAGA complex have been well-established ^5,6^, although it is less clear why some of the promoters are TFIID dependent and others are SAGA dependent. In contrast, the SAGA complex was found to play a post initiation role in metazoans^7^. Nevertheless, compared with the molecular details of ordered PIC assembly, the regulation of PIC assembly by TFs and epigenetic regulators is less well understood.

In metazoans, elongation by Pol II includes two steps, i.e. promoter-proximal pause release of Pol II (PPPRP) and productive elongation, and PPPRP and initiation are recognized as critical checkpoints of transcriptional regulation ^8^. The binding of NELF and DSIF to elongating Pol II 20 to 80 nt downstream of transcription start sites (TSSs) stabilizes its promoter-proximal pause, and the release requires kinase activity of P-TEFb, a heterodimer of CDK9 and Cyclin T1, and the PAF1 complex (PAF1C) ^9,10^. P-TEFb activity is elaborately regulate through incorporation into both the 7SK snRNP ribonucleoprotein complex, in which its kinase activity is constrained ^11^, and the multiprotein super elongation complex (SEC), in which it is active. The SEC is composed of AFF1/AFF4, AF9/ENL and ELL1 in addition to P-TEFb ^12^, and the subunits separated by a “slash” are homologous and mutually exclusive in the complex ^13^. Among the SEC subunits, AFF1, AFF4, AF9, ENL, and ELL1 are common fusion partners of MLL1 in mixed lineage leukemias (MLLs), which account for over 70% of the infant leukemia cases and approximately 10% of the adult acute myeloid leukemia (AML) cases ^14^. MLL1 is a member of the SET1/MLL family methyltransferases in human cells that affect chromatin structure and gene expression by methylation of H3K4 at key regulatory regions of the genome ^15^. Particularly, promoter-associated H3K4 trimethylation (H3K4me3) has been shown to stimulate transcriptional initiation by serving as a binding site for the TAF3 subunit of TFIID ^16^. MLL fusion proteins (MLL-FPs), arising from chromosomal translocations that lead to in-frame fusions of an *MLL1* 5’ fragment to one of the more than sixty fusion partner genes, are sufficient to induce MLL, but usually require DOT1L for the activation of key target genes with incompletely understood mechanisms ^17^.

DOT1L, the only known H3K79 methyltransferase in human cells and a homolog of yeast Dot1, normally forms a complex with AF10, AF17 and AF9/ENL, two subunits shared with SEC ^18,19^. Within DOT1L complex, DOT1L is known to antagonize deacetylases SIRT1-mediated epigenetic silencing with incompletely understood mechanisms ^20^, AF10 stimulates the conversion of H3K79 mono-methylation to di- and tri-methylation ^21^, AF9 and ENL are capable of binding acetylated H3 and facilitating the recruitment of DOT1L ^22,23^, and the function of AF17 is less well understood. Moreover, AF10 and AF17 are also fusion partners of MLL1 in MLL ^14^. However, mainly due to the association with Pol II ^24^, sharing subunits with SEC and the association of H3K79 di- and tri-methylation with the bodies of active genes, DOT1L is believed to regulate transcriptional elongation without sufficient evidence. Furthermore, the binding of Dot1/Dot1L to H2BK120 monoubiquitination (H2Bub1) is required for H3K79 methylation in both yeast and metazoans ^25^, Dot1 is known to promote H2Bub1 by inhibiting the recruitment of SAGA complex in yeast ^26^, and DOT-1.1 is known to suppress H2Bub1 in *C. elegans*^27^, but the effect and the underlying mechanisms in human cells are undetermined.

To understand roles of DOT1L complex in transcriptional regulation and MLL, we performed functional genomic studies in human cells, and discovered that DOT1L complex regulates transcriptional initiation likely by facilitating the recruitment of TFIID in both non-MLL and MLL cells and that DOT1L stimulates H2Bub1 by limiting the chromatin occupancy of human SAGA (hSAGA) complex.

## RESULTS

### DOT1L promotes the chromatin association of Pol II in human cells

We chose human erythroleukemia cell lines, HEL and K562, for understating roles of DOT1L complex in transcriptional regulation in non-MLL cells because DOT1L is required for erythropoiesis ^28^ and it is easy to perform functional genomic studies with them. Promoter-associated Pol IIs and gene body-associated Pol IIs are highly phosphorylated on serine 5 (ser-5) and serine 2 (ser-2) of their C-terminal domain (CTD), respectively, and changes of CTD phosphorylation usually reflect transcriptional changes. To examine the effects of DOT1L loss on transcription, we knocked it down by a lentiviral shRNA in HEL cells, and analyzed the effects on two forms of CTD phosphorylated Pol II by Western blot (WB). DOT1L knockdown (KD) markedly reduced the level of CTD ser-2 phosphorylation of Pol II, indicating effects on transcriptional elongation (Figure S1A). However, chromatin occupancy changes of proteins do not always follow their abundance changes. To examine the effects on Pol II chromatin occupancy, we performed ChIP-qPCR for total, ser-5 phosphorylated and ser-2 phosphorylated Pol II on *c-MYC* and *CTNNB1.* Consistent with the WB result, the chromatin occupancy of ser-2 phosphorylated Pol II decreased after DOT1L KD (Figure S1B and C). Interestingly, the chromatin occupancies of total and ser-5 phosphorylated Pol II also decreased, suggesting that the effects of DOT1L loss on transcription is likely unlimited to elongation (Figure S1D and E). To determine if DOT1L regulates global Pol II occupancy in HEL cells, we performed ChIP-seq experiments for total and ser-2 phosphorylated Pol II. DOT1L KD reduced their global occupancies, pointing to a general role of DOT1L in promoting the chromatin association of Pol II in human cells (Figure 1A, B, and C). To assess the effect of DOT1L KD on transcriptome, we performed RNA-seq experiments. A subset of genes showed expression changes with 259 downregulated and 266 upregulated after DOT1L KD (Figure S2A and B).

**Figure 1.**
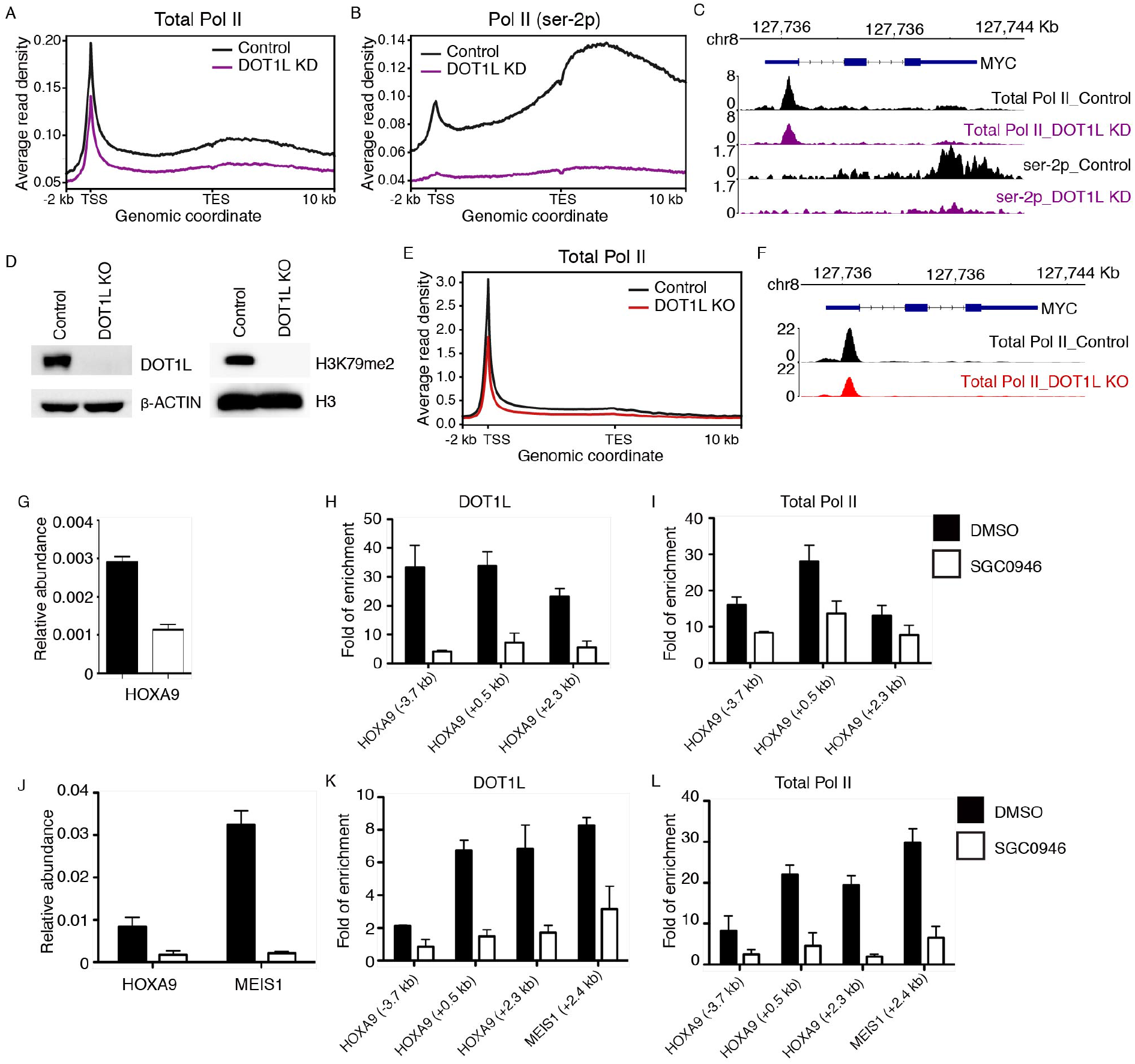
DOT1L promotes the chromatin association of Pol II in human cells. (A) and (B) Comparison of the occupancies of total Pol II (A) and Pol II (ser-2p) (B) on an average gene in DOT1L KD versus control HEL cells by ChIP-seq. (C) Normalized read distribution of total and ser2-phophorylated Pol II ChIP-seq experiments within the *c-MYC* locus in DOT1L KD versus control HEL cells. (D) Characterization of a DOT1L KO K562 cell line by Western blot. (E) Comparison of total Pol II occupancies on an average gene in control and DOT1L KO cells. (F) Normalized read distribution of total Pol II ChIP-seq within the *c-MYC* locus in DOT1L KO versus control K562 cells. (G) Comparison of the mRNA level of *HOXA9* in DMSO and SGC0946 treated THP1 cells by qRT-PCR. (H) and (I) Comparison of DOT1L (H) and total Pol II (I) occupancies within the *HOXA9* locus in DMSO and SGC0946 treated THP1 cells by ChIP followed by qPCR. (J) Comparison of the mRNA levels of *HOXA9* and *MEIS1* in DMSO and SGC0946 treated MOLM-13 cells by qRT-PCR. (K) and (L) Comparison of DOT1L (K) and total Pol II (L) occupancies within the *HOXA9* and *MEIS1* loci in DMSO and SGC0946 treated MOLM-13 cells by ChIP followed by qPCR.

To determine if DOT1L plays a general role in promoting the chromatin association of Pol II in non-MLL human cells, we performed total Pol II ChIP-seq experiments in control and DOT1L knockout (KO) K562 cells generated by the CRISPR-Cas9 technique (Figure 1D). The reduced global occupancy of Pol II after DOT1L KO supported a general role (Figure 1E and F). To determine if this is also the case in human MLL cells, we treated THP1 and MOLM-13 cells, respectively, with a potent DOT1L inhibitor, SGC0946. DOT1L inhibition reduced both the expression of and the occupancy of Pol II on common key target genes of DOT1L and MLL-AF9 (Figure 1G, H, I, J, K, and L). Altogether, these data suggested that DOT1L is a general regulator for promoting the chromatin occupancy of Pol II in non-MLL and MLL human cells.

### DOT1L may not play a major role in transcriptional elongation in human cells

Transcriptional elongation is divided into PPPRP and productive elongation. DOT1L is considered an elongation factor without sufficient evidence, and a recent study revealed that it does not regulate PPPRP in mouse ES cells, but may affect downstream elongation ^29^. To assess if it affects PPPRP in human cells, we calculated traveling ratio (TR) ^30^ of Pol II with total Pol II ChIP-seq data from HEL and K562 cells, respectively (Figure 2A). DOT1L KD or KO had little effect on the TR of Pol II (Figure 2B and C), consistent with results of the earlier study that DOT1L may not play a major role in PPPRP in human cells. To further validate this finding, we performed PRO-seq in control and DOT1L KD HEL cells to analyze the distribution of engaged Pol II and calculate TR. DOT1L KD reduced the chromatin occupancy but had almost no effect on the TR of engaged Pol II (Figure 2D, E and F), further supporting that DOT1L is unlikely to play a major role in the regulation of PPPRP. To determine if DOT1L affects the rate of productive elongation, we performed 4sUDRB-seq experiments, which is based on the reversible inhibition of transcriptional elongation with DRB and the labeling of newly transcribed RNA with uridine analog 4-thiouridine (4sU) ^31^. We found that DOT1L KO minimally affected the productive elongation rate of Pol II (Figure 2G, H, I, J and K). Altogether, these data suggested that DOT1L may not play a major role in transcriptional elongation in human cells.

**Figure 2.**
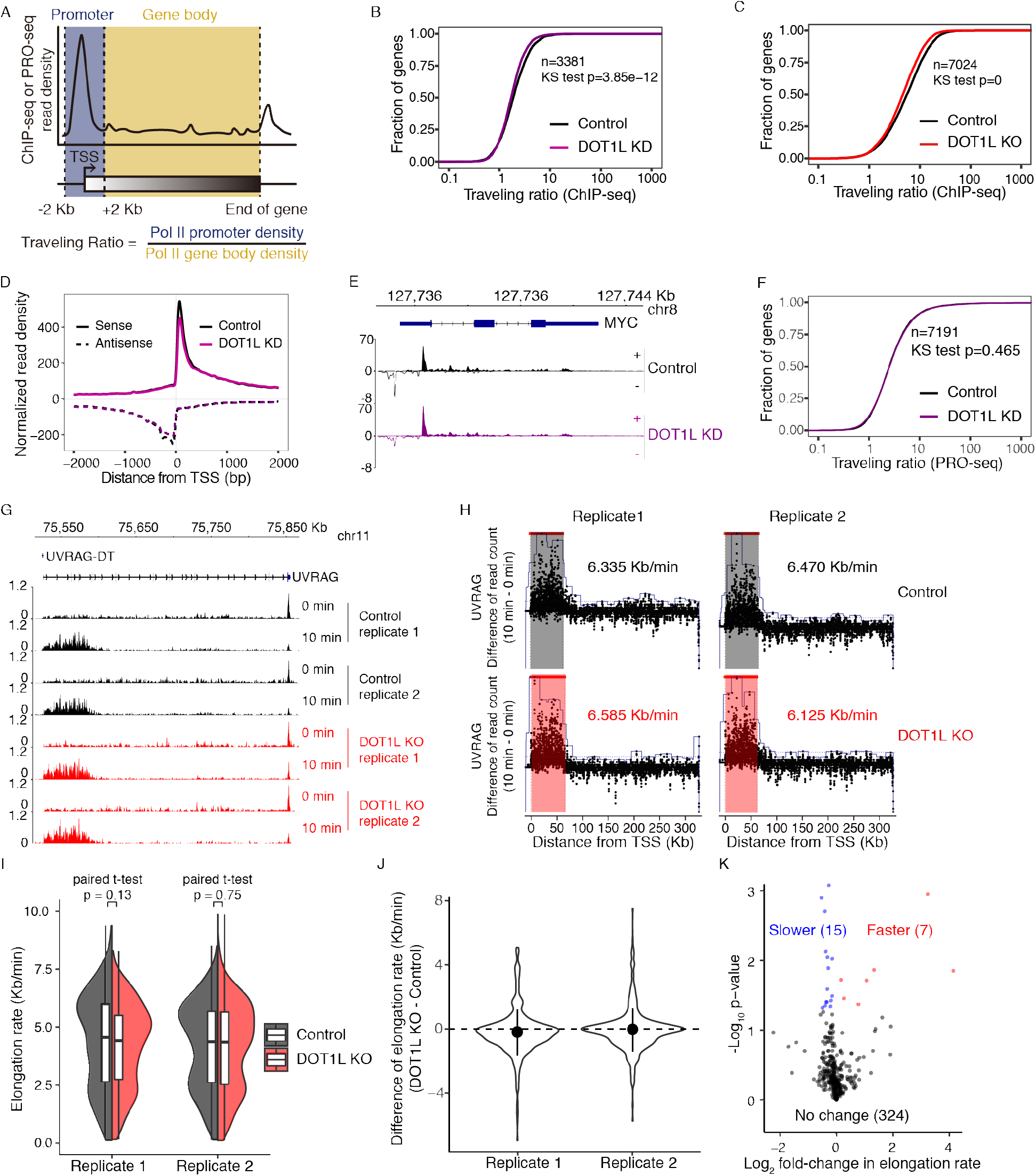
DOT1L may not play a major role in transcriptional elongation in human cells. (A) Schematic representation describing the calculation used to determine the traveling ratio (TR) at each Pol II-bound gene. (B) Comparison of the TR of total Pol II in control and DOT1L KD HEL cells. (C) Comparison of the TR of total Pol II in control and DOT1L KO K562 cells. (D) Comparison of the distribution of engaged Pol II near TSSs in control and DOT1L KD HEL cells by PRO-seq. (E) Normalized read distribution of PRO-seq within the *c-MYC* locus in DOT1L KD versus control HEL cells. (F) Comparison of the TR of engaged Pol II in control and DOT1L KD HEL cells. (G) Normalized read distribution of 4sUDRB-seq experiments comparing Pol II elongation rate of *UVRAG* in DOT1L KO versus control K562 cells. (H) HMM model of elongation rate calculation for *UCRAG* in control and DOT1L KO K562 cells. (I) Range of Pol II elongation rate on genes in control and DOT1L KO K562 cells. (J) Pol II elongation rate change in DOT1L KO versus control K562 cells. (K) A volcano plot of genes with Pol II elongation rate changes in DOT1L KO versus control K562 cells.

### DOT1L regulates transcriptional initiation in human cells

Ordered binding of TBP, TFIIA and TFIIB to promoters precedes and facilitates Pol II recruitment in transcriptional initiation in eukaryotic cells. Reduced Pol II occupancy near TSSs in DOT1L depleted cells raised the possibility that DOT1L may regulate transcriptional initiation. To test if that is the case, we performed ChIP-seq experiments for TBP, TFIIA and TFIIB in control and DOT1L KO cells. DOT1L KO markedly reduced the global occupancies of TBP and TFIIA and the occupancy of TFIIB on a subset of genes (Figure 3A, B, C, D, E, F and G), suggesting that DOT1L play a general role in the regulation of transcriptional initiation.

**Figure 3.**
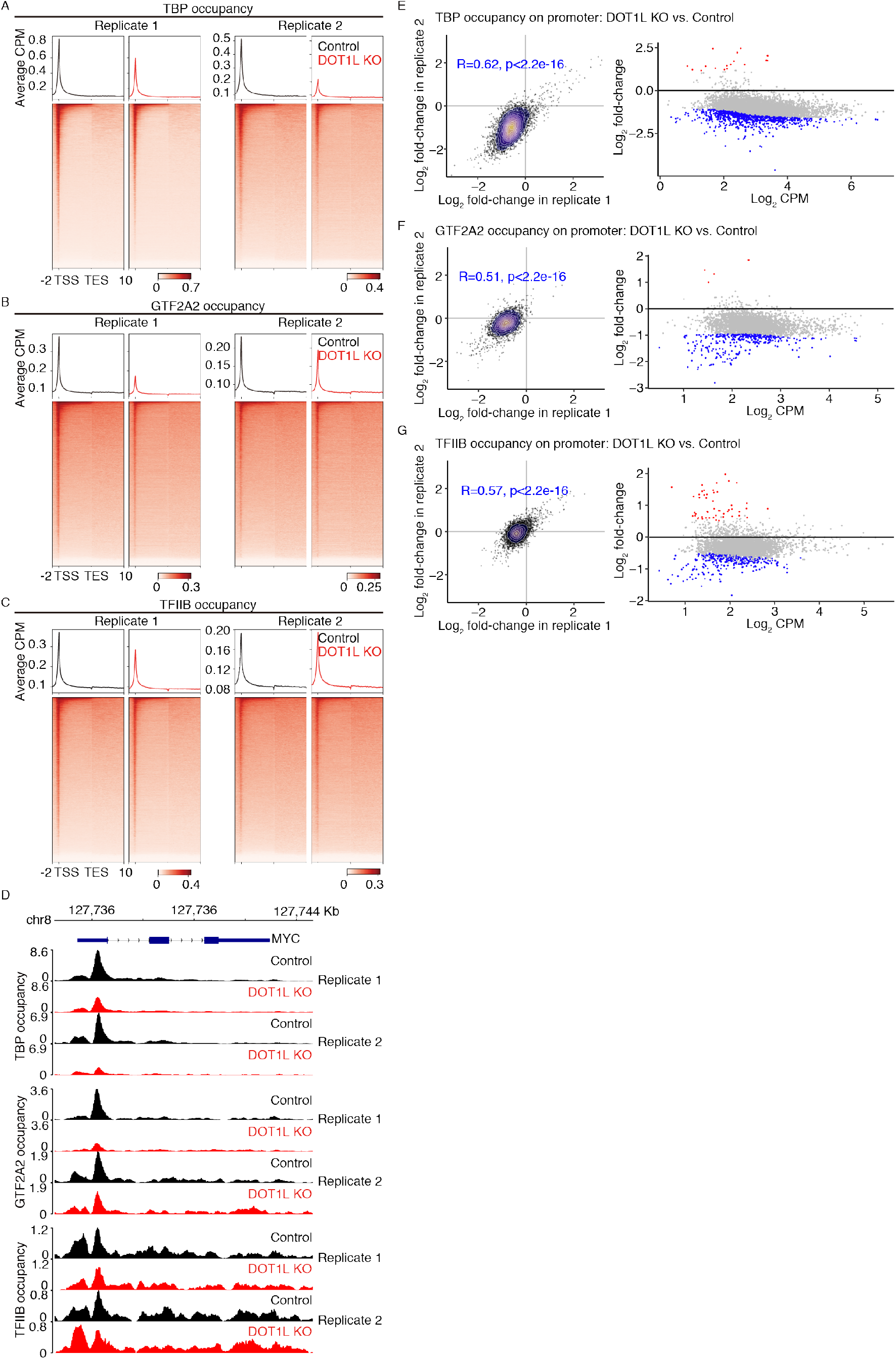
DOT1L regulates transcriptional initiation in human cells. (A), (B) and (C) Genome-wide meta-gene profiles and heatmaps of ChIP-seq comparing the chromatin occupancies of TBP (A), TFIIA (B) and TFIIB (C) in DOT1L KO versus control K562 cells. (D) Normalized read distribution of ChIP-seq experiments comparing the occupancy of TBP, TFIIA, and TFIIB within the *c-MYC* locus in DOT1L KO versus control K562 cells. (E), (F) and (G) Occupancy changes of TBP (E), TFIIA (F) and TFIIB (G) on promoters. Left panels, Dot and density plots of occupancy changes on promoters in two replicates. Consistency between the replicates was measured by Pearson correlation coefficient. Right panels, MA plots of differential occupancy on promoters based on the replicates.

### DOT1L recruits TFIID via physical interactions

Mechanistically, DOT1L may regulate transcriptional initiation via physical interactions with initiation factors or creating binding sites for them by methylating H3K79. Pulldown assays using synthesized short H3 fragments harboring methylated K79 to identify binders of H3K79 methylation were known to be unspecific because of the hydrophobic nature of the fragments. We therefore chose to understand mechanisms underlying DOT1L-mediated transcriptional initiation by unbiased analysis of its interacting proteins. To this end, we performed large-scale co-immunoprecipitation (co-IP) using a DOT1L antibody with nuclear extract from K562 cells, and characterized immunoprecipitated proteins by mass spectrometry (Figure 4A). Besides subunits of DOT1L complex, we also identified several TAFs, TFIIH, hSAGA complex, Mediator, etc. which were previously unknown to interact with DOT1L (Figure 4B). The identification of TAFs raised a possibility that DOT1L may regulate the recruitment of TFIID via physical interactions. The confirmation of the interaction between DOT1L and TAF7 by co-IP experiments from both directions suggested that likely to be the case (Figure 4C and D). We were also able to confirm the interaction between DOT1L complex and Mediator by co-IP experiments from both directions, further supporting a role of DOT1L in transcriptional initiation (Figure 4E and F).

**Figure 4.**
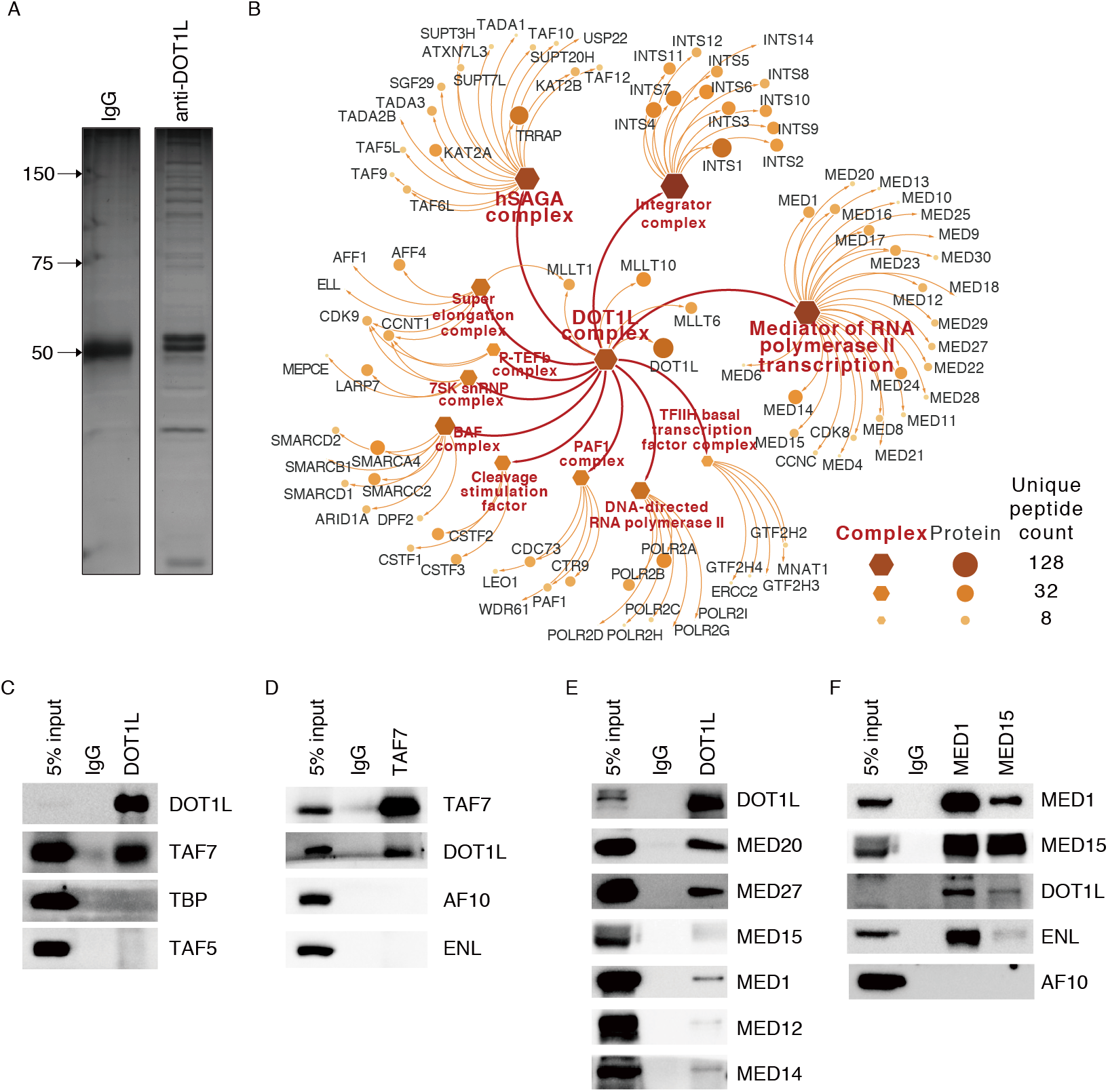
DOT1L interacts with transcriptional initiation factors. (A) Silver stain of proteins immunoprecipitated by rabbit IgG and a DOT1L antibody, respectively, and separated on an SDS-PAGE gel. (B) Protein complexes co-immunoprecipitated with DOT1L. (C) WB analyses of TFIID subunits co-immunoprecipitated with DOT1L. (D) WB analyses of DOT1L complex subunits co-immunoprecipitated with TAF7. (E) WB analyses of Mediator subunits co-immunoprecipitated with DOT1L. (F) WB analyses of subunits of DOT1L complex co-immunoprecipitated with MED1 and MED15.

### ENL regulates transcriptional initiation in human cells

ENL/AF9 are shared subunits between DOT1L complex and SEC, and are capable of recruiting the two complexes to chromatin by binding to acetylated H3 through theirs YEATS domains ^22,23^. In addition, ENL has been shown to affect the chromatin occupancy of Pol II and regulate PPPRP as one of the subunits of SEC ^23^. Our discovery of DOT1L as a general regulator of transcriptional initiation in this study and a previously discovered role of ENL in DOT1L recruitment raised a possibility that ENL may also regulate transcriptional initiation. To test this idea, we performed ChIP-seq experiments for TBP in control and ENL KO K562 cells generated by the CRISPR-Cas9 technique (Figure 5A). The loss of ENL markedly reduced the global occupancy of TBP (Figure 5B, C, D and E), supporting a general role in the regulation transcriptional initiation. The next question we asked was if ENL regulates productive elongation. To test this idea, we performed 4sUDRB-seq experiments in control and ENL KO cells, and found that ENL KO minimally affected the productive elongation rate of Pol II (Figure 5F, G, H, I and J). Altogether, these data suggested that in addition to a recognized role in the regulation of PPPRP, ENL also regulates transcription initiation but may not a play a major role in productive elongation.

**Figure 5.**
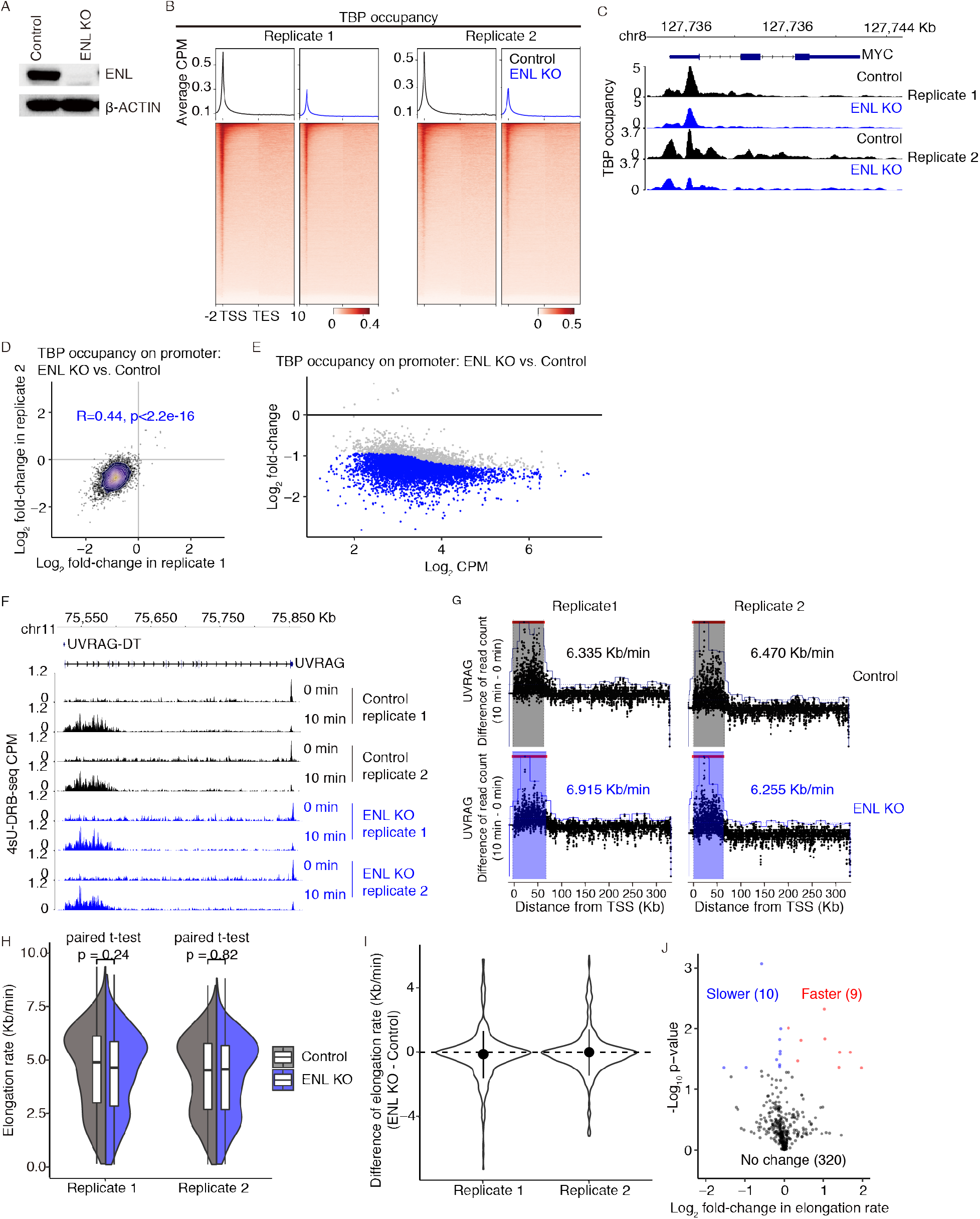
ENL regulates transcriptional initiation in human cells. (A) Characterization of an ENL KO K562 cell line by Western blot. (B) Genome-wide meta-gene profiles and heatmaps of ChIP-seq comparing the chromatin occupancies of TBP in ENL KO versus control K562 cells. (C) Normalized read distribution of ChIP-seq comparing the occupancy of TBP within the *c-MYC* locus in ENL KO versus control K562 cells. (D) A dot and density plot of TBP occupancy change on promoters in two replicates. Consistency between the replicates was measured by Pearson correlation coefficient. (E) An MA plot of differential occupancy on promoters calculated with the replicates. (F) Normalized read distribution of 4sUDRB-seq experiments comparing the Pol II elongation rate of *UVRAG* in ENL KO versus control K562 cells. (G) HMM model of Pol II elongation rate calculation for *UCRAG* in control and ENL KO K562 cells. (H) Range of Pol II elongation rate in control and ENL KO K562 cells. (I) Pol II elongation rate changes in ENL KO versus control K562 cells. (J) A volcano plot of genes with Pol II elongation rate changes in ENL KO versus control K562 cells.

### Neither DOT1L nor ENL affects global chromatin accessibility

H3K79 methylation is globally associated with active genes and a conformation change of H3 is required for the methylation of K79 by DOT1L ^25^, which raised a possibility that DOT1L may regulate global chromatin accessibility and therefore transcriptional initiation. To test this idea, we perform ATAC-seq in control, DOT1L KO and ENL KO K562 cells. We found that the DOT1L KO slightly increased global chromatin accessibility but ENL KO had no effect on it (Figure 6A and B). A closer examination of the peaks uncovered that DOT1L KO slightly decreased the percentage of peaks on promoters, but ENL KO exhibited little effect (Figure 6C). However, further analyses revealed no global reduction of promoter accessibility after DOT1L or ENL KO but an expected positive correlation between promoter accessibility and mRNA level change (Figure 6D). Altogether, these data suggested that although H3K79 methylation is associated with active genes, neither DOT1L nor ENL affects global chromatin accessibility.

**Figure 6.**
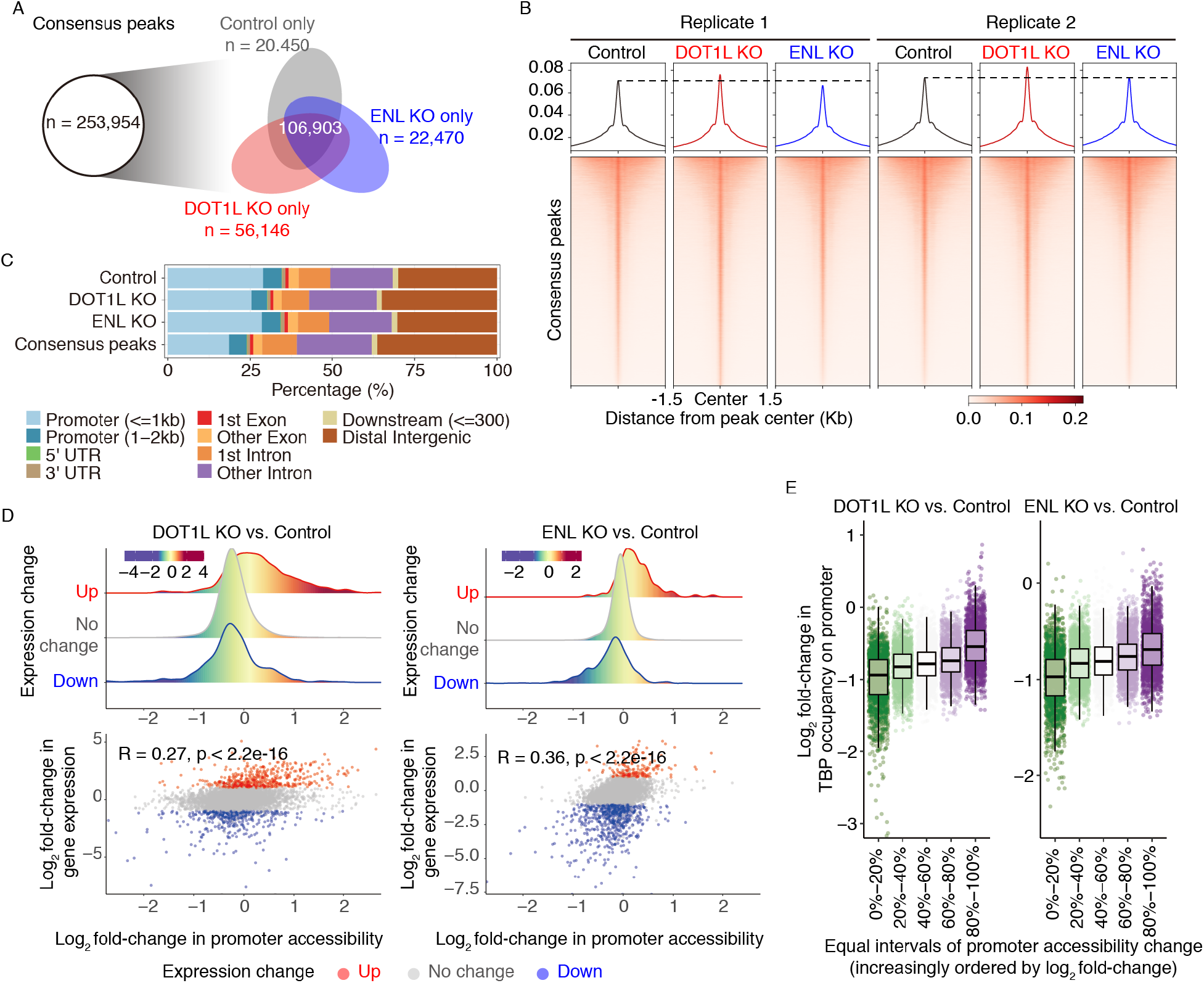
Neither DOT1L nor ENL affects global chromatin accessibility. (A) Venn diagram showing overlaps among peaks identified from control, DOT1L KO and ENL KO cells, respectively. The consensus peak set was obtained by merging peaks from all the cell lines. (B) Chromatin accessibility around consensus peak centers in control, DOT1L KO and ENL KO cells. (C) Genomic feature annotation of ATAC-seq peaks identified in control, DOT1L KO cells, ENL KO cells and the consensus peak set. (D) Promoter accessibility changes of differentially expressed genes in control, DOT1L KO and ENL KO cells. Top, Promoter accessibility changes of up-regulated, down-regulated and genes without significant change. Bottom, Scatter plots of gene expression change versus promoter accessibility change. Pearson’s correlation coefficient was labeled on top left. (E) Box and jitter plots of TBP occupancy change on promoters grouped by degree of accessibility changes in control, DOT1L KO and ENL KO cells.

To further understand the relationships among DOT1L KO, TBP occupancy and promoter accessibility, we divided promoters into five groups according to their values of accessibility changes after DOT1L KO and compared their TBP occupancy changes. As expected, TBP occupancy was found to be positively corelated with promoter accessibility (Figure 6E). Notably, we also found that even promoters with increased accessibility exhibited decreased occupancies of TBP after DOT1L KO (Figure 6E), strongly supporting a DOT1L dependency of TFIID recruitment regardless of promoter accessibility change.

### DOT1L promotes H2Bub1 by limiting the recruitment of hSAGA complex

In both yeast and metazoans, H2Bub1 is required for H3K79 methylation by Dot1. SAGA complex is known to catalyze H3 acetylation, in particular H3K9 and H3K27 acetylation, and H2B deubiquitination. In yeast, Dot1 was found to promote H2Bub1 by limiting the recruitment of SAGA complex in an enzymatic activity independent manner ^26^, in *C. elegans,* DOT-1.1 was found to suppress H2Bub1 ^27^, but the effect and the underlying mechanisms in human cells are undetermined. The identification of human SAGA (hSAGA) complex in our DOT1L immunoprecipitants led us to study this matter. The interaction between DOT1L and hSAGA complex was confirmed by co-IP experiments from both directions (Figure 7A and B). To determine if DOT1L affects the recruitment of hSAGA complex and H2Bub1, we performed ChIP-seq experiments for PCAF, a subunit of the acetyltransferase module of hSAGA complex, and H2Bub1 in control and DOT1L KO K562 cells. Consistent with the results in yeast, the loss of DOT1L increased the chromatin occupancy of PCAF but decreased global H2Bub1 (Figure 7C, D, and E). We noted that the quality of the PCAF ChIP-seq was not high. To confirm the results, we performed CUT&Tag for PCAF in control and DOT1L KO cells, and got the same results (Figure 7F and G). Taken together, these data suggested that in human cells, DOT1L promote H2Bub1 by limiting the recruitment of hSAGA complex likely through physical interactions.

**Figure 7.**
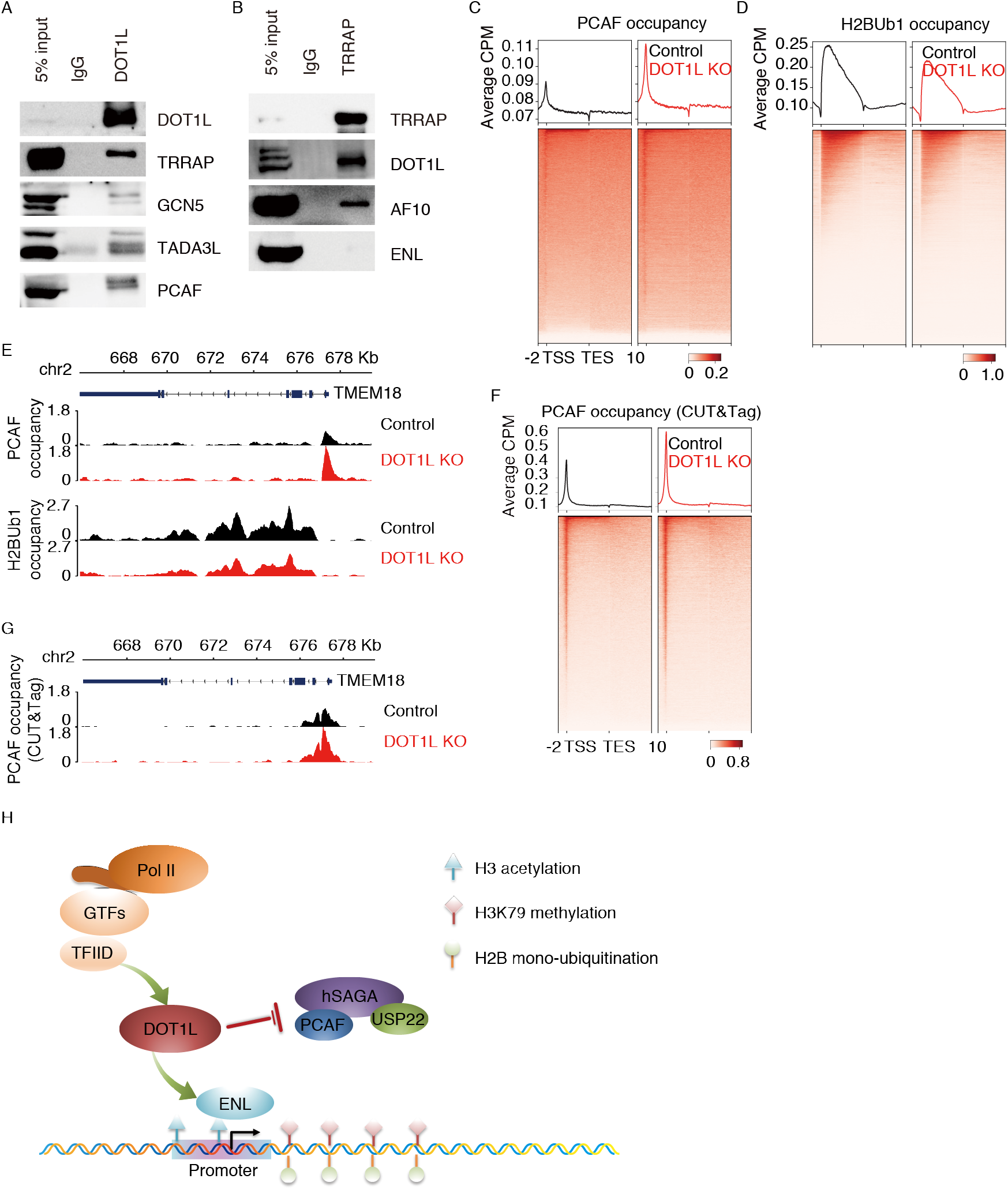
DOT1L promotes H2Bub1 by limiting the recruitment of hSAGA complex. (A) WB analyses of subunits of hSAGA complex co-immunoprecipitated with DOT1L. (B) WB analyses of subunits of DOT1L complex co-immunoprecipitated with TRRAP. (C) and (D) Genome-wide meta-gene profiles and heatmaps of ChIP-seq comparing the occupancies of PCAF (C) and H2Bub1 (D) in DOT1L KO versus control K562 cells. (E) Normalized read distribution of ChIP-seq comparing the occupancy of PCAF and H2Bub1 within the *TMEM18* locus in DOT1L KO versus control K562 cells. (F) Genome-wide meta-gene profiles and heatmaps of CUT&Tag comparing the occupancies of PCAF in DOT1L KO versus control K562 cells. (G) Normalized read distribution of CUT&Tag comparing the occupancy of PCAF within the *TMEM18* locus in DOT1L KO versus control K562 cells. (H) Working model of this study. The ENL subunit of DOT1L complex binds acetylated H3 and recruits DOT1L, DOT1L complex regulates transcriptional initiation likely by recruiting TFIID. DOT1L promotes H2Bub1 by limiting the recruitment of hSAGA complex, which contains a deubiqutinase module.

## DISCUSSION

DOT1L normally forms a complex with AF10, AF17 and ENL/AF9, is considered an elongation factor without sufficient evidence, and is dysregulated in most of the cases of MLL with incompletely understood mechanisms. In this study, we provide results suggesting that DOT1L complex actually regulates transcriptional initiation by facilitating the recruitment of TFIID and that DOT1L promotes H2Bub1 by limiting the recruitment of hSAGA complex (Figure 7H).

### DOT1L complex regulates transcriptional initiation

DOT1L was considered an elongation factor for association with Pol II, sharing ENL/AF9 subunits with SEC, a critical elongation complex, and the localization of H3K79me2 on the bodies of active genes. By analyzing both traveling ratio and elongation rate of Pol II, we found that DOT1L is unlikely to play a major role in PPPRP and productive elongation in human cells with the conclusion related to PPPRP in agreement with that of a recent published study using mouse ES cells ^29^. By analyzing GTFs binding and their interactions with DOT1L, we found that DOT1L actually regulates global transcriptional initiation likely through physical interactions with GTFs. In addition, we found that ENL, one of the shared subunits of SEC and DOT1L complex that have been reported to be able to recruit DOT1L by binding acetylated H3 ^23^, also regulates global transcriptional initiation but not productive elongation. Altogether, our results defined DOT1L complex as a general regulator of transcriptional initiation although the molecular details need further investigation.

### DOT1L stimulates H2Bub1 by limiting the recruitment of hSAGA complex

In both yeast and metazoans, H2Bub1 is required for H3K79 methylation by providing binding site for Dot1/Dot1L. In yeast, Dot1 was found to promote H2Bub1 by limiting the recruitment of SAGA complex in an enzymatic activity independent manner ^26^, in *C. elegans,* DOT-1.1 was found to suppress H2Bub1 ^27^, but the effect and the underlying mechanisms in human cells were undetermined. In the yeast, the role of SAGA complex in transcriptional initiation has been well-established ^5,6^, but it is less clear why some of the promoters are TFIID dependent and others are SAGA dependent. In contrast, the SAGA complex was found to play a post initiation role in metazoans^7^. We found that DOT1L promotes H2Bub1 by limiting the recruitment of hSAGA complex, which not only is in agreement with the conclusion in yeast but also suggests that the connection between the two complexes is likely to be evolutionarily conserved. Our results also raised a possibility that Dot1 may facilitate the choose between TFIID and SAGA complex for transcriptional initiation at least in yeast. Nevertheless, the functional significance and the molecular details of this connection remain to be elucidated.

### MLL-FPs promote transcriptional initiation through DOT1L complex

MLL-FPs are known to induce MLL by maintaining the expression of several key target genes, most notably *HOXA9* and *MESI1,* which normally are highly expressed in hematopoietic progenitor cells for promoting proliferation and protection from stress ^32,33^. Considering that major fusion partners of MLL1 are subunits of either SEC or DOT1L complex, that SEC is a critical elongation factor and DOT1L was considered an elongation factor, MLL-FPs were believed to achieve so by stimulating transcriptional elongation. Our discovery of DOT1L complex as a general regulator of transcriptional initiation suggests that MLL-FPs are likely able to stimulate transcriptional initiation and elongation. The binding to CpG islands near promoters via the CXXC domain within their MLL1 N-terminus fragments and the stimulation of transcriptional initiation and elongation by recruiting DOT1L complex and P-TEFb, respectively, enable MLL-FPs to efficiently maintain the expression of key target genes.

## METHODS

### Cells and cell culture

Human cells HEL, THP1, and MOLM-13 cells were cultured in RPMI-1640 + 10% FBS + 2% Penicillin/Streptomycin + 2 mM L-Glutamine + 55 μM ß-mercaptoethanol. Human cells K562 were cultured in 90% DMEM + 10% FBS+ 2% Penicillin/Streptomycin + 2 mM L-Glutamine.

### CUT&Tag and data analyses

CUT&Tag experiments were performed as previously described with minor modifications ^34^. Briefly, 100,000 cells were used for each experiment. Cells were bound to Concanavalin A-coated beads without fixation and chromatin opening. After primary and secondary antibodies binding, pA-Tn5 transposome binding, and tagmentation, DNA was extracted and amplified by PCR. Raw reads were filtered using fastp ^35^ (version 0.13.1, default parameters) and aligned using Bowtie2 ^36^ (version 2.3.4.1) to Bowtie2 index based on hg38 downloaded from NCBI. Low-quality alignments were filtered out using SAMtools ^37^ (version 0.1.19) with command “samtools view -F 1804 -q 25”. MarkDuplicates tools in Picard ^38^ was used to identify and remove PCR duplicates from the aligned reads. We used bamCoverage from deepTools ^39^ (version 3.3.1) to calculate read coverage per 50-bp bin using the CPM normalization option. Heatmaps were generated using computeMatrix and plotHeatmap, and meta-gene profile plots were generated using computeMatrix and plotProfile from deepTools.

### Accession numbers

Next generation sequencing data have been submitted to GEO repository under accession number GSE161367.

## ACKNOWLEDGEMENTS

The authors would like to thank H. Jiang (UVA) for critical reading of the manuscript. R.G.R. is supported by a Leukemia and Lymphoma Society SCOR grant, and M.Y. is supported by grants from The Program for Professor of Special Appointment (Eastern Scholar) at Shanghai Institutions of Higher Learning, and National Natural Science Foundation of China (31671351).

## AUTHOR CONTRIBUTIONS

M.Y. and R.G.R. designed the experiments, J.Z., T.T., L.C., M.Y., Z.L., and L.F. performed the experiments and analyzed the data, A.W. performed the bioinformatics analysis, and M.Y., A.W. and R.G.R wrote the paper.

## SUPPLEMENTARY INFORMATION

### SUPPLEMENTARY FIGURES

**Figure S1.**
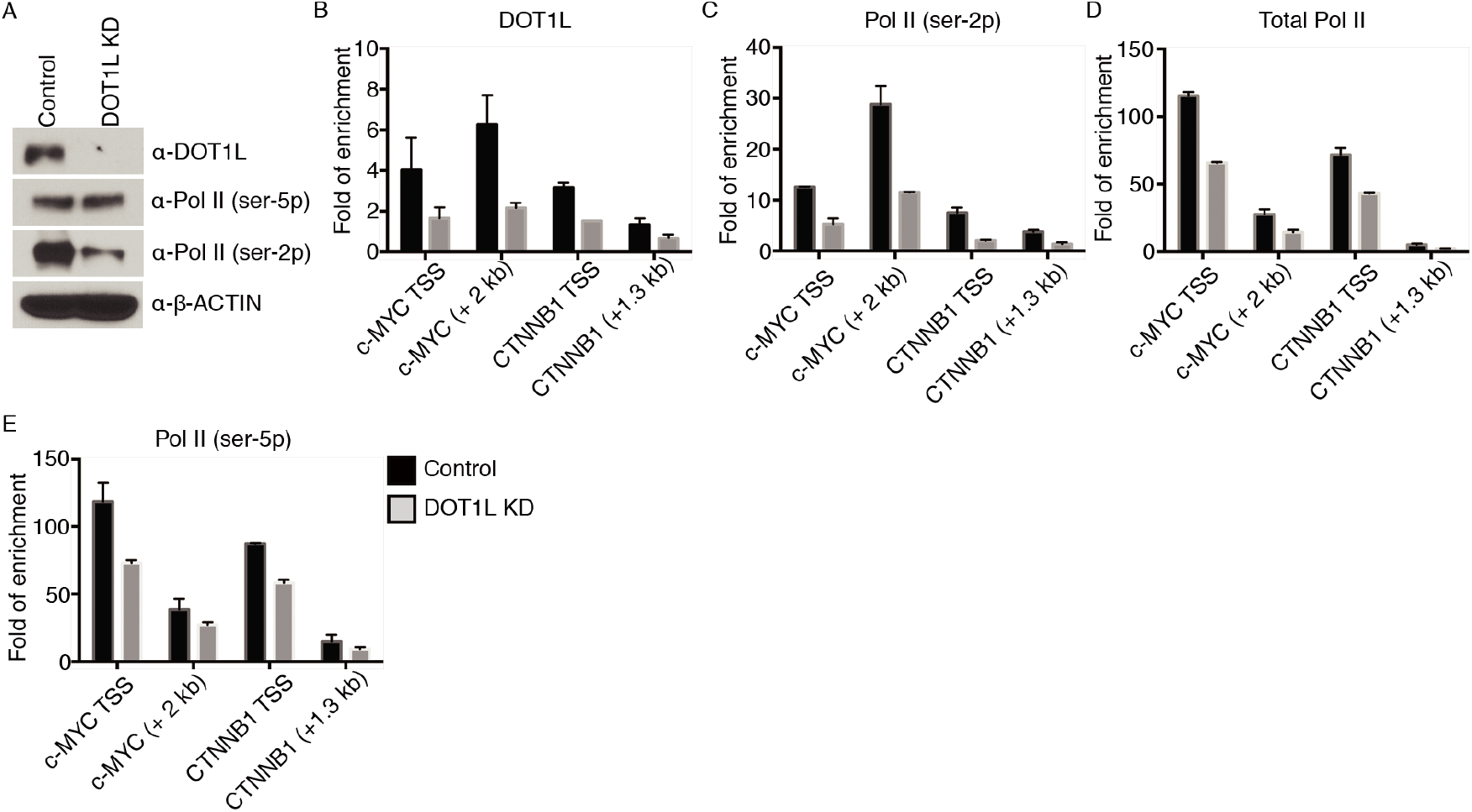
DOT1L KD reduced the chromatin occupancy of Pol II in HEL cells. (A) Comparison of the levels of DOT1L, Pol II (ser-5p) and Pol II (ser-2p) in control versus DOT1L KD HEL cells by Western blot. β-ACTIN was used as a loading control. (B), (C), (D), and (E) Comparison of the occupancies of DOT1L (B), Pol II (ser-2p) (C), total Pol II (D), and Pol II (ser-5p) (E) on *c-MYC* and *CTNNB1* by ChIP-qPCR in control versus DOT1L KD HEL cells.

**Figure S2.**
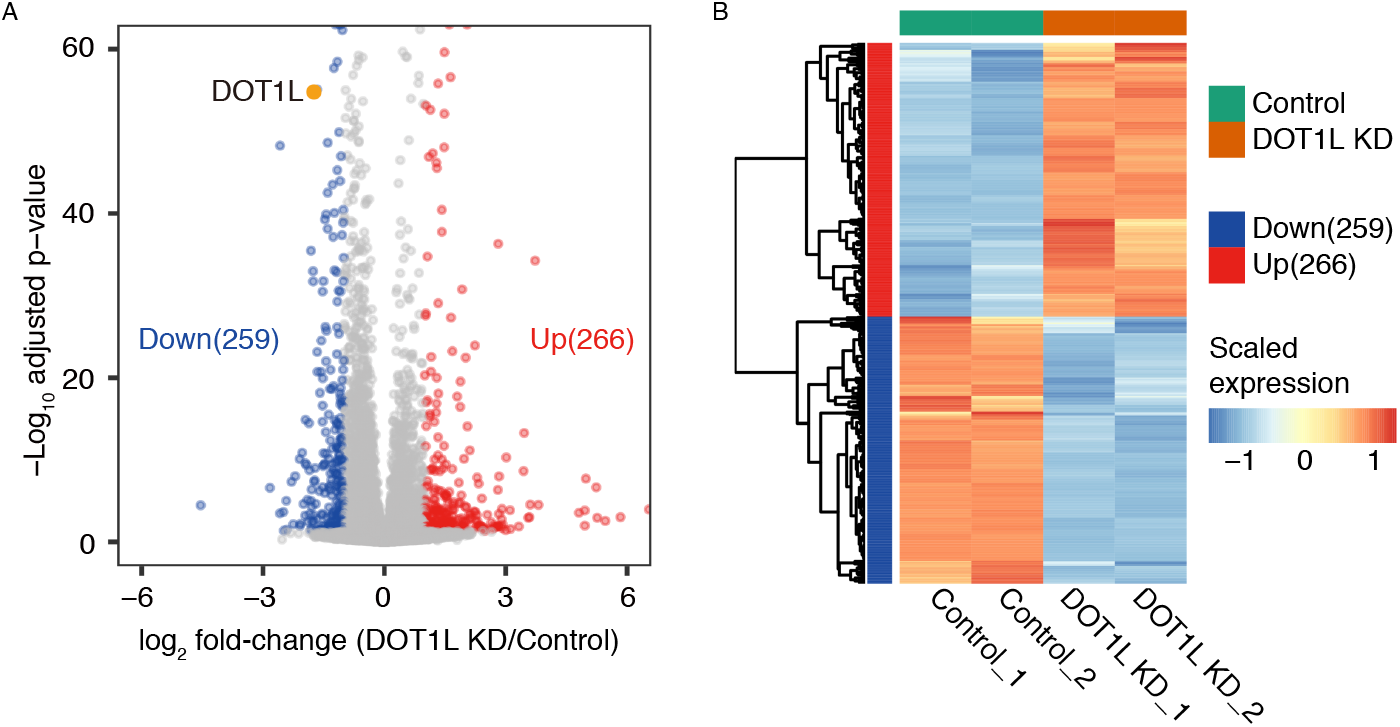
DOT1L KD affected the transcription of a subset of genes in HEL cells. (A) A volcano plot of differentially expressed genes in control and DOT1L KD HEL cells. (B) A heatmap of differentially expressed genes in control and DOT1L KD HEL cells.

**Figure S3.**
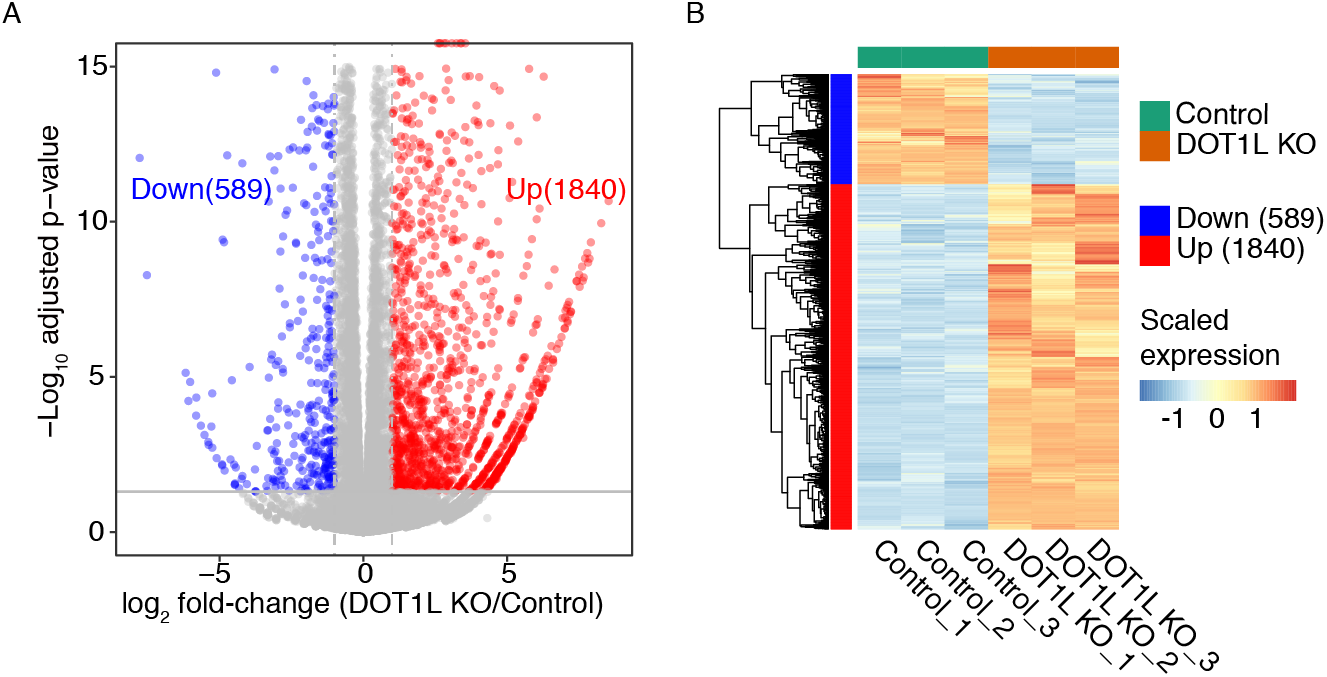
DOT1L KO affected the transcription of a subset of genes in K562 cells. (A) A volcano plot of differentially expressed genes in control and DOT1L KO K562 cells. (B) A heatmap of differentially expressed genes in control and DOT1L KO K562 cells.

### SUPPLEMENTARY TABLES

**Table S1.**
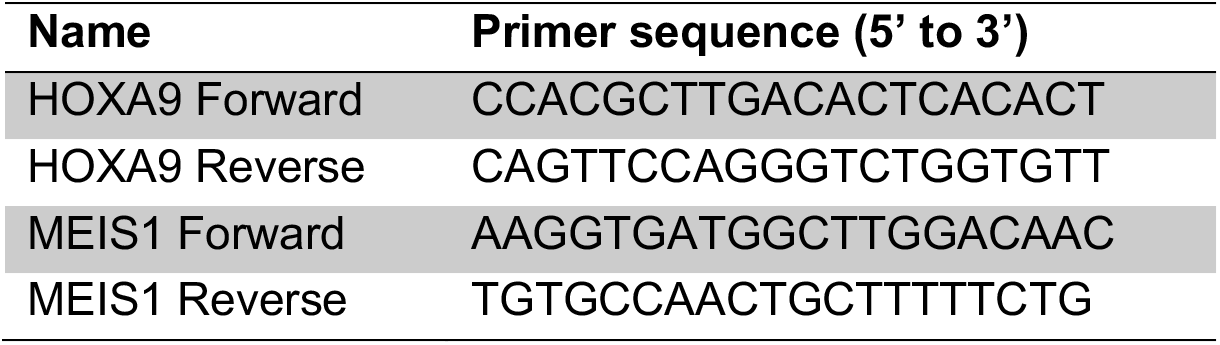
Primers for qRT-PCR.

**Table S2.**
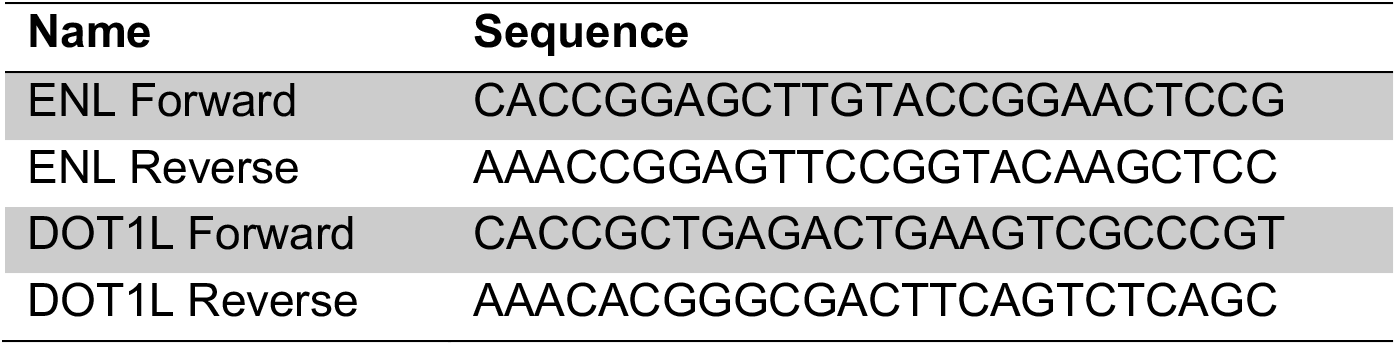
Guide RNAs.

**Table S3.** Antibodies.

| Target | Vender | Catalogue number | Applications |
| --- | --- | --- | --- |
| DOT1L | Bethyl | A300-953A | WB |
| DOT1L | Cell Signaling | 77087S | WB, IP |
| Pol II | Santa Cruz | sc-899 | WB, ChIP |
| ser-2 phosphorylated Pol II | Active Motif | 61083 | WB |
| ser-5 phosphorylated Pol II | Active Motif | 61085 | WB |
| TBP | Santa Cruz | sc-56795 | WB, ChIP |
| GTF2A2 | Proteintech | 10540-1-AP | WB, ChIP |
| TFIIB | Proteintech | 16467-1-AP | WB, ChIP |
| H3K79me2 | Abcam | ab3594 | WB |
| TAF7 | Proteintech | 13506-1-AP | WB, IP |
| MED1 | Bethyl | A300-793A | WB, IP |
| MED27 | Santa Cruz | sc-390296 | WB |
| MED15 | Proteintech | 11566-1-AP | WB, IP |
| MED12 | Proteintech | 20028-1-AP | WB |
| MED14 | ABclonal | A10455 | WB |
| MED20 | ABclonal | A15757 | WB |
| ENL | Santa Cruz | sc-393196 | WB |
| TRRAP | Abcam | ab183517 | WB, IP |
| TADA3L | Proteintech | 10839-1-AP | WB |
| H2Bub1 | Active Motif | 39623 | ChIP |
| PCAF | Abcam | ab12188 | ChIP, CUT&Tag |
| AF10 | Santa Cruz | sc-53156 | WB |
| GCN5 | ABclonal | A2224 | WB |

**Table S4.**
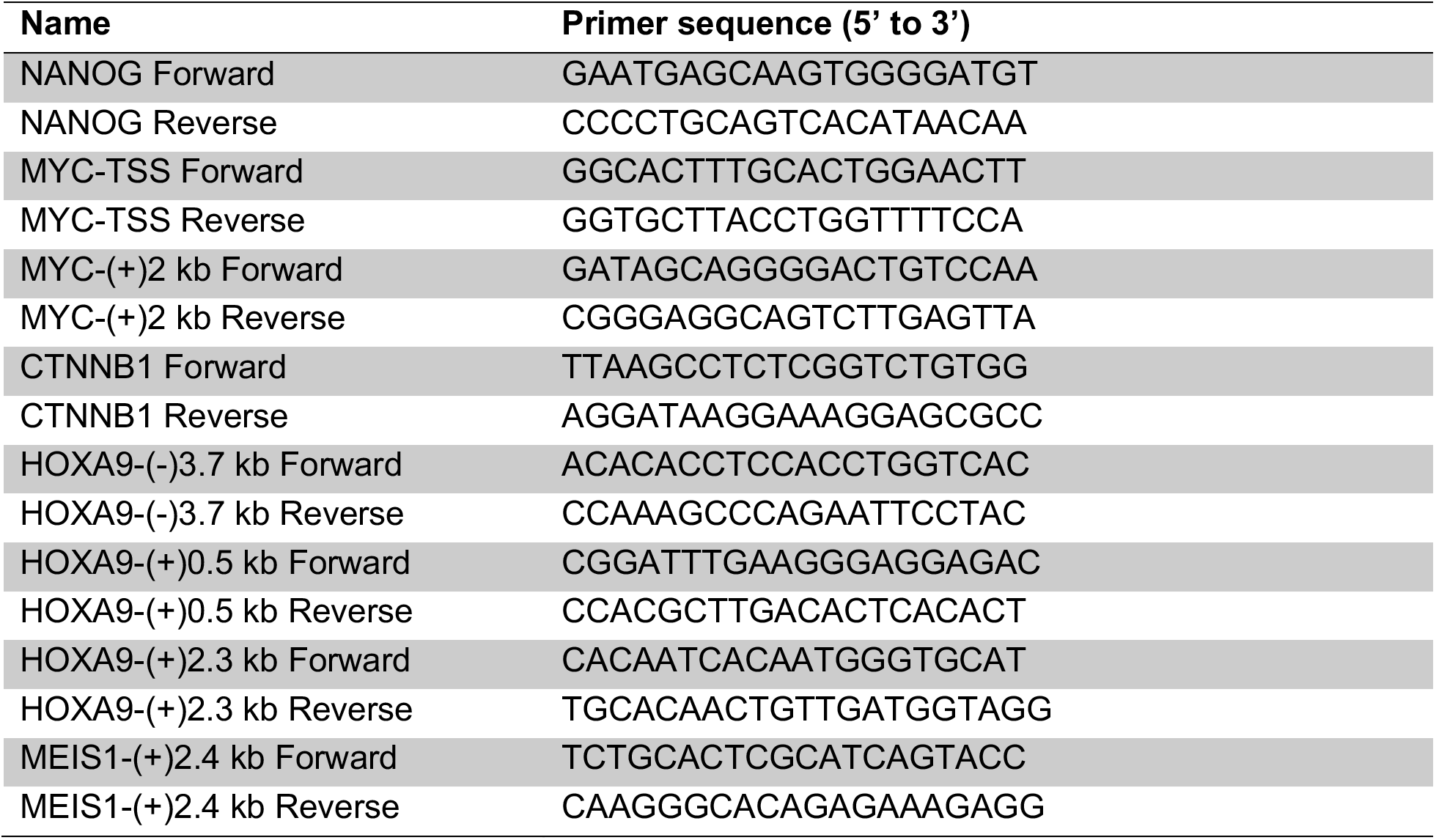
Primers for ChIP-qPCR.

### SUPPLEMENTARY METHODS

#### RNA interference, RNA extraction, reverse transcription, real-time PCR, RNA-seq and data analyses

pLKO.1 TRC control and mission shRNA clones were purchased from Sigma Aldrich. For lentivirus production and transduction, 60-90% confluent 293T cells in antibiotic-free medium were transfected on day 1 with TRC control or gene-specific shRNA clones along with packaging plasmids psPAX2 and pMD2.G. On day 2 in the morning, medium containing transfection reagent was replaced by fresh medium containing 2% Penicillin-Streptomycin (Sigma Aldrich, cat. no. P0781). On day 3 in the afternoon, cells were resuspended in virus-containing medium, and spun at 2,000 rpm at 20 °C for 1 hour. After spin infection, virus-containing medium was removed and cells were resuspended in fresh medium and cultured overnight. On day 4 in the morning, cells were washed twice with PBS and resuspended in fresh medium. On day 5 in the morning, puromycin was added to a final concentration of 2 μg/ml. Cells were cultured for additional 72 hours before being harvested for qRT-PCR, Western blot, and ChIP. Mission shRNA clone and qRT-PCR primers used in this study are listed in Table S1.

*RNA was extracted from cells using RNeasy Plus Mini Kit (Qiagen, cat. no. 74134) and Quick-RNA MiniPrep Kit (Zymo Research, cat. no. R1054) by following the manufacturers’ protocols. Libraries of strand-specific RNA-seq were constructed as previously described^40^. Raw reads were filtered using fastp^35^ (version 0.13.1, default parameters) and mapped to hg38 using HISAT2 ^41^ (version 2.1.0) with parameters “--rna-strandness RF -dta”. Read counts per gene were calculated in strand-specific manner using featureCounts^42^. For each RNA-seq library, reads were mapped with overall mapping rate of ~97%. Differential expression analysis were performed using DESeq2 ^43^, and genes with adjusted p value < 0.05 and fold change of > 2 were identified as significantly differentially expressed.*

#### Generation of knockout cell lines by CRISPR/Cas9

A K562-derived cell line inducibly expressing Cas9, K562-iCas9, was generated by transducing pCW-Cas9-Hygro into K562 cells and selecting clones with high-level expression. Guide RNAs (gRNAs) were designed using the tool provided by Benching, cloned into lentiGuide-Puro, individually transduced into K562-iCas9 cells. Single colonies obtained by serial dilution were expanded and subsequently characterized by Western blot. Sequences of gRNAs used in this study are listed in Table S2.

#### ChIP, ChIP-sequencing (ChIP-seq) and data analysis

ChIP assays were performed as previously described ^44^. Normally, cells were fixed with 0.4% (v/v) formaldehyde at room temperature for 10 min. To improve the ChIP efficiency, double fixation was used. For double fixation with EGS (Thermo, cat. no. 21565) and formaldehyde, cells were fixed initially with 1.5 mM EGS at room temperature for 30 min, and subsequently with 0.4% formaldehyde at room temperature for 10 min. For double fixation with DMA (Thermo, cat. no. 20660) and formaldehyde, cells were fixed initially with 25 mM DMA at room temperature for 1 hour, and subsequently with 0.4% formaldehyde at room temperature for 10 min. For sonication, fixed cells were washed twice with PBS and resuspended in ice-cold RIPA-0.3 buffer (10 mM Tris-HCl, 1 mM EDTA, 1% Triton X-100, 0.1% SDS, 0.1% NaDOC, and 0.3 M NaCl, pH 7.4) supplemented with Protease Inhibitor Cocktail (Millipore, cat. no. 535140) at a concentration of 40 million cells/ml; genomic DNA was disrupted to a size range of 100 to 500 bp. For immunoprecipitation, on day 1 antibodies were diluted in RIPA-0.3 and bound to Dynabeads protein A (Thermo, cat. no. 10002D) by incubating at 4 °C for 3 hours. Afterwards, the bead-antibody complexes were washed twice with RIPA-0.3 and then incubated with sonicated chromatin at 4 °C overnight. On day 2, after 2 washes with RIPA-0.5, 1 wash with RIPA-0.3, 1 wash with RIPA-0, 2 washes with LiCl buffer (10 mM Tris-HCl, 1 mM EDTA, 0.25 M LiCl, 0.25% NP-40, and 0.25% NaDOC, pH 7.4), and 2 washes with TE buffer, bound protein-DNA complexes were resuspended in elution buffer (10 mM Tris-HCl, 1mM EDTA, 0.2 M NaCl, and 1% SDS, pH 7.4) supplemented with 10 μg/ml RNase A for both elution and RNA digestion, and incubated at 55 °C for 1 hour. Then Proteinase K was added to a final concentration of 200 μg/ml, and after 30 min incubation the temperature was increased to 65 °C for crosslink reversal. After incubation for 4 to 6 hours, DNA was purified by ChIP DNA Clean & Concentrator (Zymo Research, cat. no. D5205). For qPCR, a site 2 kb downstream of the TSS of the *NANOG* gene was used as an internal control, and fold-enrichment was calculated by the 2^ΔCt^ method. The antibodies that were used for ChIP assays are listed in Table S3. Real-time PCR primers for ChIP are listed in Table S4.

ChIP-seq libraries were constructed with 2–10 ng immunoprecipitated DNA. After end-repair, A-tailing, and barcode ligation, barcoded DNA was amplified by 16- to 18-cycle PCR. Libraries were sequenced on Illumina HiSeq 2000, HiSeq 2500 or HiSeq X Ten by following the manufacturer’s protocols. Raw reads were filtered using fastp ^35^ (version 0.13.1, default parameters) and aligned using Bowtie2 ^36^ (version 2.3.4.1) to Bowtie2 index based on hg38 downloaded from NCBI. Low-quality alignments were filtered out using SAMtools ^37^ (version 0.1.19) with command “samtools view -F 1804 -q 25”. MarkDuplicates tools in Picard ^38^ was used to identify and remove PCR duplicates from the aligned reads. Peak calling was carried out using MACS2 ^45^ version 2.2.6 with input control. Broad peaks were called with parameters “- q 0.01 --broad --nomodel --shift 0 --keep-dup all”. We used bamCoverage from deepTools ^39^ (version 3.3.1) to calculate read coverage per 50-bp bin using the CPM normalization option. Heatmaps were generated using computeMatrix and plotHeatmap, and meta-gene profile plots were generated using computeMatrix and plotProfile from deepTools. To visualize promoter occupancy change of GTFs in MA plot, read counts in promoters overlap with corresponding broad peaks were used as input of edgeR ^46^, and log2 CPM and log2 fold-change were estimated with norm.factors set to be proportional to total mapped read count after removing duplications. For each gene, promoter and gene body read densities were calculated as read count normalized by length and sequencing depth, and traveling ratio was calculated as promoter read density divided by gene body read density for Pol II-bound genes, which were defined by CPM in promoter larger than 4.

#### PRO-seq and data analyses

PPO-seq experiments were performed as previously described ^47^, and the libraries were sequenced by Illumina HiSeq 2500. Raw reads were filtered using fastp ^35^ (version 0.13.1, default parameters) and aligned using Bowtie2 ^36^ (version 2.3.4.1) to Bowtie2 index based on hg38 downloaded from NCBI. Low-quality alignments were filtered out using SAMtools ^37^ (version 0.1.19) with command “samtools view -F 4 -q 10”. Strand-specific meta-gene profiles were generated using the groHMM package ^48^ from Bioconductor. Strand-specific read coverage was calculated using “bedtools genomecov” with “-ibam”, “-bg” and “-strand” options and normalized by total mapped reads before loading to IGV for visualization.

#### 4sUDRB-seq and data analyses

4sUDRB-seq experiments were performed as previously described ^31^ with minor modifications. Briefly, 10 million cells were used for each experiment. After DRB (Sigma, cat. no. D1916) treatment and 4sU (Sigma, cat. no. T4509) incorporation, total RNA was extracted. After RNA biotinylation and free biotin removal, biotinylated RNA was purified by streptavidin-coupled Dynabeads (Thermo, cat. no. 11205D). Before library construction, rRNA was depleted by following a published protocol ^49^. Sequencing libraries were constructed by following the

Illumina TruSeq RNA Library preparation protocol. 4sU-DRB-seq reads were filtered using fastp ^35^ (version 0.13.1, default parameters) and aligned to the human genome hg19 using Bowtie2 ^36^ (version 2.3.4.1) with parameter “-N 1”. Only paired reads aligned to the same chromosome, not to chromosome chrUn_gl000220 (rRNA) and with alignment scores < 5 were kept using awk. rRNA percentage was calculated for each sample. Average rRNA percentage is 3.1% for 0 min samples and 0.24% for 10 min. BamCoverage from deepTools ^39^ (version 3.3.1) was used to generate bigwig files of normalized read coverage per 50-bp bin.

Transcripts were filtered to calculate elongation rate. For each gene, the longest transcript was chosen, and it was required to have a minimum transcript length of 30 kb and do not contain other transcription start site (TSS) of this gene. Further filtering was performed to exclude transcripts overlapping with other genes or within 2 kb from TSSs of another genes. Finally, 3707 transcripts were kept for calculating elongation rates. Advancing waves were identified using a three state Hidden Markov Model (HMM) that was previously developed and implemented on GRO-seq data from a human cell line ^50^, in which model, 2kb regions around TES were not included for the unstable signal in them. Paired t-test was used to test whether the elongation rate distribution of DOT1L or ENL KO cells is significantly different from that of control cells for each replicate respectively. Significant faster- or slower elongating genes was identified using Welch’s t-test based on both two replicates of each cell line.

#### ATAC-seq and data analyses

Tn5 transposase expression and purification, and transposome assembly was conducted as previously described ^51^. ATAC-seq experiments were performed by following a published protocol ^52^. Briefly, 50,000 cells were used for each experiment. After nuclei preparation, tagmentation, termination and DNA purification, samples were amplified by PCR with one universal forward primer and different indexed reverse primers. ATAC-seq pair-end reads were filtered using fastp ^35^ (version 0.13.1, default parameters) and aligned to the human genome hg38 using Bowtie2 ^36^ (version 2.3.4.1) with parameter “-X 2000”. Samtools was used to filter for reads mapped to Chr1-22 and ChrX, and MarkDuplicates tools in Picard ^38^ was used to identify and remove PCR duplicates from the aligned reads. The final deduplicated BAM file was used in the downstream analyses.

Tn5 transposase insertions, which refer to the precise single-base locations where Tn5 transposase accessed the chromatin, were identified by correcting the read start positions by a constant offset (“+” stranded +4 bp, “-” stranded -5 bp). To generate depth-normalized accessibility track, bigwig files were constructed based on the Tn5 offset-corrected insertion sites using GenomicRanges ^53^ and rtracklayer ^54^ packages in R. Meta-gene profile plots were generated using computeMatrix and plotProfile from deepTools ^39^. For each replicate, peak calling was performed on the Tn5-corrected single-base insertions using the “MACS2 callpeak” command with parameters “-g hs -n Ctrl -q 0.01 --shift −19 --extsize 38 --nomodel --nolambda --keep-dup all --call-summits”. The peaks were then filtered to remove peaks overlapping hg38 blacklisted region (http://mitra.stanford.edu/kundaje/akundaje/release/blacklists/hg38-human/hg38.blacklist.bed.gz). Consistent peaks of each cell type were called on pooled replicates, followed by filtering for those displaying at least 50% overlap with any peak from each of the two single replicate peak sets. Consensus peak set was obtained by merge the consistent peaks identified in each cell type using command “bedtools merge”. Peak annotation of consistent peaks of each cell type and the final consensus peak set was performed using ChIPseeker^55^ package in Bioconductor.

#### Co-immunoprecipitation (co-IP), large scale co-IP, and mass spectrometry

Co-IP assays were performed as previously described with minor modification ^56^. Nuclear extract (NE) from HEL cells was diluted 2 to 3 fold by adding NE dilution buffer. Antibodies were incubated with Dynabeads protein A at 4 °C for 3 hours, and then cross-linked to beads by 25 mM DMA (Pierce, cat. no. 20660) at room temperature for 1 hour. Normally, 0.5 to 1 mg nuclear extract was used for each co-IP. After overnight incubation at 4 °C, bead-antibody-protein complexes were washed with BC-150 buffer (10 mM Tris-HCl, 0.2 mM EDTA, 150 mM KCl, 20% glycerol, and 0.1% NP-40, pH 7.9) for 4 times. The proteins were eluted from beads by 50 mM glycine (pH 2.4), immediately neutralized in 1 M Tris-HCl, pH 7.4, and analyzed by Western blot. For large-scale co-IP, 10 μg antibody and ~75 mg NE were used for each experiment. Bead-antibody-protein complexes were washed with BC-200 buffer (10 mM Tris-HCl, 0.2 mM EDTA, 200 mM KCl, 20% glycerol, and 0.1% NP-40, pH 7.9) for 6 times followed by with 1 x PBS twice. Eluted proteins were analyzed by a Multidimensional Protein Identification Technology (MudPIT) system.

MS/MS spectra were searched using MASCOT engine (Matrix Science, London, UK; version 2.2) against the Human UniProt database (downloaded at May 2019; 171145 sequences). For protein identification, the following options were used: Peptide mass tolerance=20 ppm, MS/MS tolerance=0.1 Da, Enzyme=Trypsin, Missed cleavage=2, Fixed modification: Carbamidomethyl (C), Variable modification: Oxidation(M); score > 20. In total, 1980 and 1960 non-redundant protein were identified in DOT1L-IP and IgG-IP samples, respectively. Proteins identified in DOT1L-IP sample were filtered by: 1) Unique peptide count of a protein in IgG-IP sample (UPC_IgG_) is no larger than 1; 2) If UPC_IgG_ of a protein is 0, unique peptide count of the same protein in DOT1L-IP sample (UPC_DOT1L_) should be at least 2, and if UPC_IgG_ of a protein is 1, UPC_DOT1L_ should be at least 3. 677 proteins met the requirement were referred to as DOT1L-accosiated proteins. To identify candidate DOT1L-associated complexes, 2916 human protein complexes were downloaded from CORUM (version 3.0), which is a comprehensive resource of mammalian protein complexes, and 157 redundant ones with duplicated “ComplexName” and “subunits UniProt ID” and 89 complexes containing only one subunit were filtered out. Out of the remaining 2670 complexes, coverage of 63 were no less than 70%, and 41/63 were further filtered out for being totally contained by one of the remaining 22 complexes. After manual review and correction, 11 complexes, including DOT1L complex itself, were considered as reliable DOT1L-associated complexes.

## REFERECES

1. Bhagwat, A.S. & Vakoc, C.R. Targeting Transcription Factors in Cancer. Trends Cancer 1, 53–65 (2015).

2. Roeder, R.G. 50+ years of eukaryotic transcription: an expanding universe of factors and mechanisms. Nat Struct Mol Biol 26, 783–791 (2019).

3. Nogales, E., Louder, R.K. & He, Y. Structural Insights into the Eukaryotic Transcription Initiation Machinery. Annu Rev Biophys 46, 59–83 (2017).

4. Koutelou, E., Hirsch, C.L. & Dent, S.Y. Multiple faces of the SAGA complex. Curr Opin Cell Biol 22, 374–82 (2010).

5. Donczew, R., Warfield, L., Pacheco, D., Erijman, A. & Hahn, S. Two roles for the yeast transcription coactivator SAGA and a set of genes redundantly regulated by TFIID and SAGA. Elife 9(2020).

6. Helmlinger, D. & Tora, L. Sharing the SAGA. Trends Biochem Sci 42, 850–861 (2017).

7. Weake, V.M. et al. Post-transcription initiation function of the ubiquitous SAGA complex in tissue-specific gene activation. Genes Dev 25, 1499–509 (2011).

8. Adelman, K. & Lis, J.T. Promoter-proximal pausing of RNA polymerase II: emerging roles in metazoans. Nat Rev Genet 13, 720–31 (2012).

9. Vos, S.M. et al. Structure of activated transcription complex Pol II-DSIF-PAF-SPT6. Nature 560, 607–612 (2018).

10. Yu, M. et al. RNA polymerase II-associated factor 1 regulates the release and phosphorylation of paused RNA polymerase II. Science 350, 1383–6 (2015).

11. Peterlin, B.M. & Price, D.H. Controlling the elongation phase of transcription with P-TEFb. Mol Cell 23, 297–305 (2006).

12. Smith, E., Lin, C. & Shilatifard, A. The super elongation complex (SEC) and MLL in development and disease. Genes Dev 25, 661–72 (2011).

13. Lu, H. et al. Gene target specificity of the Super Elongation Complex (SEC) family: how HIV-1 Tat employs selected SEC members to activate viral transcription. Nucleic Acids Res 43, 5868–79 (2015).

14. Krivtsov, A.V. & Armstrong, S.A. MLL translocations, histone modifications and leukaemia stem-cell development. Nat Rev Cancer 7, 823–33 (2007).

15. Ernst, P. & Vakoc, C.R. WRAD: enabler of the SET1-family of H3K4 methyltransferases. Brief Funct Genomics 11, 217–26 (2012).

16. Lauberth, S.M. et al. H3K4me3 interactions with TAF3 regulate preinitiation complex assembly and selective gene activation. Cell 152, 1021–36 (2013).

17. Bernt, K.M. et al. MLL-rearranged leukemia is dependent on aberrant H3K79 methylation by DOT1L. Cancer Cell 20, 66–78 (2011).

18. Wang, X., Chen, C.W. & Armstrong, S.A. The role of DOT1L in the maintenance of leukemia gene expression. Curr Opin Genet Dev 36, 68–72 (2016).

19. Wood, K., Tellier, M. & Murphy, S. DOT1L and H3K79 Methylation in Transcription and Genomic Stability. Biomolecules 8(2018).

20. Chen, C.W. et al. DOT1L inhibits SIRT1-mediated epigenetic silencing to maintain leukemic gene expression in MLL-rearranged leukemia. Nat Med 21, 335–43 (2015).

21. Deshpande, A.J. et al. AF10 regulates progressive H3K79 methylation and HOX gene expression in diverse AML subtypes. Cancer Cell 26, 896–908 (2014).

22. Li, Y. et al. AF9 YEATS domain links histone acetylation to DOT1L-mediated H3K79 methylation. Cell 159, 558–71 (2014).

23. Wan, L. et al. ENL links histone acetylation to oncogenic gene expression in acute myeloid leukaemia. Nature 543, 265–269 (2017).

24. Kim, S.K. et al. Human histone H3K79 methyltransferase DOT1L protein [corrected] binds actively transcribing RNA polymerase II to regulate gene expression. J Biol Chem 287, 39698–709 (2012).

25. Worden, E.J. & Wolberger, C. Activation and regulation of H2B-Ubiquitin-dependent histone methyltransferases. Curr Opin Struct Biol 59, 98–106 (2019).

26. van Welsem, T. et al. Dot1 promotes H2B ubiquitination by a methyltransferase-independent mechanism. Nucleic Acids Res 46, 11251–11261 (2018).

27. Cecere, G., Hoersch, S., Jensen, M.B., Dixit, S. & Grishok, A. The ZFP-1(AF10)/DOT-1 complex opposes H2B ubiquitination to reduce Pol II transcription. Mol Cell 50, 894–907 (2013).

28. Feng, Y. et al. Early mammalian erythropoiesis requires the Dot1L methyltransferase. Blood 116, 4483–91 (2010).

29. Cao, K. et al. DOT1L-controlled cell-fate determination and transcription elongation are independent of H3K79 methylation. Proc Natl Acad Sci USA (2020).

30. Rahl, P.B. et al. c-Myc regulates transcriptional pause release. Cell 141, 432–45 (2010).

31. Fuchs, G. et al. 4sUDRB-seq: measuring genomewide transcriptional elongation rates and initiation frequencies within cells. Genome Biol 15, R69 (2014).

32. Unnisa, Z. et al. Meis1 preserves hematopoietic stem cells in mice by limiting oxidative stress. Blood 120, 4973–81 (2012).

33. Lawrence, H.J. et al. Loss of expression of the Hoxa-9 homeobox gene impairs the proliferation and repopulating ability of hematopoietic stem cells. Blood 106, 3988–94 (2005).

34. Kaya-Okur, H.S. et al. CUT&Tag for efficient epigenomic profiling of small samples and single cells. Nat Commun 10, 1930 (2019).

35. Chen, S., Zhou, Y., Chen, Y. & Gu, J. fastp: an ultra-fast all-in-one FASTQ preprocessor. Bioinformatics 34, i884–i890 (2018).

36. Langmead, B. & Salzberg, S.L. Fast gapped-read alignment with Bowtie 2. Nat Methods 9, 357–9 (2012).

37. Li, H. et al. The Sequence Alignment/Map format and SAMtools. Bioinformatics 25, 2078–9 (2009).

38. Institute, B. Picard Toolkit. in Broad Institute, GitHub repository (2019).

39. Ramirez, F. et al. deepTools2: a next generation web server for deep-sequencing data analysis. Nucleic Acids Res 44, W160–5 (2016).

## SUPPLEMENTARY REFERENCES

6. Helmlinger, D. & Tora, L. Sharing the SAGA. Trends Biochem Sci 42, 850–861 (2017).

30. Rahl, P.B. et al. c-Myc regulates transcriptional pause release. Cell 141, 432–45 (2010).

33. Lawrence, H.J. et al. Loss of expression of the Hoxa-9 homeobox gene impairs the proliferation and repopulating ability of hematopoietic stem cells. Blood 106, 398894 (2005).

37. Li, H. et al. The Sequence Alignment/Map format and SAMtools. Bioinformatics 25, 2078–9 (2009).

38. Institute, B. Picard Toolkit. in Broad Institute, GitHub repository (2019).

40. Zhong, S. et al. High-throughput illumina strand-specific RNA sequencing library preparation. Cold Spring Harb Protoc 2011, 940–9 (2011).

41. Kim, D., Paggi, J.M., Park, C., Bennett, C. & Salzberg, S.L. Graph-based genome alignment and genotyping with HISAT2 and HISAT-genotype. Nat Biotechnol 37, 907–915 (2019).

42. Liao, Y., Smyth, G.K. & Shi, W. featureCounts: an efficient general purpose program for assigning sequence reads to genomic features. Bioinformatics 30, 923–30 (2014).

43. Love, M.I., Huber, W. & Anders, S. Moderated estimation of fold change and dispersion for RNA-seq data with DESeq2. Genome Biol 15, 550 (2014).

44. Yu, M. et al. RNA polymerase II-associated factor 1 regulates the release and phosphorylation of paused RNA polymerase II. Science 350, 1383–1386 (2015).

45. Zhang, Y. et al. Model-based analysis of ChIP-Seq (MACS). Genome Biol 9, R137 (2008).

46. McCarthy, D.J., Chen, Y. & Smyth, G.K. Differential expression analysis of multifactor RNA-Seq experiments with respect to biological variation. Nucleic Acids Res 40, 4288–97 (2012).

47. Kwak, H., Fuda, N.J., Core, L.J. & Lis, J.T. Precise maps of RNA polymerase reveal how promoters direct initiation and pausing. Science 339, 950–3 (2013).

48. Chae, M., Danko, C.G. & Kraus, W.L. groHMM: a computational tool for identifying unannotated and cell type-specific transcription units from global run-on sequencing data. BMC Bioinformatics 16, 222 (2015).

49. Morlan, J.D., Qu, K. & Sinicropi, D.V. Selective depletion of rRNA enables whole transcriptome profiling of archival fixed tissue. PLoS One 7, e42882 (2012).

50. Danko, C.G. et al. Signaling pathways differentially affect RNA polymerase II initiation, pausing, and elongation rate in cells. Mol Cell 50, 212–22 (2013).

51. Picelli, S. et al. Tn5 transposase and tagmentation procedures for massively scaled sequencing projects. Genome Res 24, 2033–40 (2014).

52. Corces, M.R. et al. An improved ATAC-seq protocol reduces background and enables interrogation of frozen tissues. Nat Methods 14, 959–962 (2017).

53. Lawrence, M. et al. Software for computing and annotating genomic ranges. PLoS Comput Biol 9, e1003118 (2013).

54. Lawrence, M., Gentleman, R. & Carey, V. rtracklayer: an R package for interfacing with genome browsers. Bioinformatics 25, 1841–2 (2009).

55. Yu, G., Wang, L.G. & He, Q.Y. ChIPseeker: an R/Bioconductor package for ChIP peak annotation, comparison and visualization. Bioinformatics 31, 2382–3 (2015).

56. Yu, M. et al. Insights into GATA-1-mediated gene activation versus repression via genome-wide chromatin occupancy analysis. Mol Cell 36, 682–95 (2009).

